# A Comparison of fMRI and Behavioral Models for Predicting Inter-Temporal Choices

**DOI:** 10.1101/866285

**Authors:** Felix G. Knorr, Philipp T. Neukam, Juliane H. Fröhner, Holger Mohr, Michael N. Smolka, Michael Marxen

## Abstract

In an inter-temporal choice (IteCh) task, subjects are offered a smaller amount of money immediately or a larger amount at a later time point. Here, we are using trial-by-trial fMRI data from 363 recording sessions and machine learning in an attempt to build a classifier that would ideally outperform established behavioral model given that it has access to brain activity specific to a single trial. Such methods could allow for future investigations of state-like factors that influence IteCh choices.

To investigate this, coefficients of a GLM with one regressor per trial were used as features for a support vector machine (SVM) in combination with a searchlight approach for feature selection and cross-validation. We then compare the results to the performance of four different behavioral models.

We found that the behavioral models reached mean accuracies of 90% and above, while the fMRI model only reached 54.84% at the best location in the brain with a spatial distribution similar to the well-known value-tracking network. This low, though significant, accuracy is in line with simulations showing that classifying based on signals with realistic correlations with subjective value produces comparable, low accuracies. These results emphasize the limitations of fMRI recordings from single events to predict human choices, especially when compared to conventional behavioral models. Better performance may be obtained with paradigms that allow the construction of miniblocks to improve the available signal-to-noise ratio.

## 1 Introduction

In intertemporal choice (ITeCh) tasks, subjects have to decide between two rewards, one being larger than the other but also paid out at a later time. To model individual behavior, it is common practice to use a hyperbolic discounting model (Kirby and Maraković, 1995; Simpson and Vuchinich, 2000). This model has a free parameter - *k* - that determines the subjective value (SV) of a delayed amount (see Fig. 2a)). It expresses the fact that we devalue things continuously the longer we have to wait for them. The *k* parameter defines how strong this effect is. Additionally, a decision likelihood function models the probability for a particular decision with an additional parameter given the SV of each offer. Using e.g. Baye’s Law (Pooseh et al., 2017), it is then possible to estimate the model parameters if the decisions are know. The additional parameter (*β* or γ, depending on the probability function) can be interpreted as a measure for decision consistency. One popular choice for this decision likelihood function is the sigmoidal Softmax function (see Fig. 2 b)) (Peters et al., 2012; Gläscher et al., 2009; Radu et al., 2011; Pine et al., 2009; Miedl, 2012). Another option is the Power model (Wulff and van den Bos, 2017) (see Methods). The fitted behavioral model parameters are often used as a subject trait. In this paper, we use them to predict the subjects decisions. We will quantify the accuracy of such predictions and compare these with predictions based on functional magnetic resonance imaging data (fMRI). As brain regions such as the ventral striatum, medial prefrontal cortex and posterior cingulate cortex have been shown to track subjective values (Kable and Glimcher, 2007), the fMRI blood oxygen level-dependent (BOLD) signal does, in principle, contain information to allow such predictions.

By nature, the behavioral models are static, that is, unlike the BOLD signal, they do not capture dynamic brain states that may contain additional, predictive information about individual choices. This is especially relevant for offers at the indifference point of a behavioral model, meaning that the probability for a particular choice is close to 50% and, thus, the expected prediction accuracy will only be at chance level. Therefore, we hypothesize that the fMRI model outperforms the behavior models. If BOLD signal-based predictions would outperform behavioral models, we would be able to use this information in the future to study dynamic influences of brain states during ITeCh choices.

For BOLD signal-based predictions, an often used multivariate algorithm is the linear Support Vector Machine (SVM) (Norman et al., 2006; Kamitani and Tong, 2005; Mitchell et al., 2004; Wang et al., 2015). It is a machine learning algorithm, that learns to map observations to classes from examples, the training data. The target classes (labels) in this case are either ‘sooner’ or ‘later’ for the two decision options. As inputs, also called feature vectors, we used voxel-wise coefficients of single-trial regressors within a univariate fMRI model from small regions-of-interest called ‘search lights’ (Soon et al., 2008; Haynes et al., 2007; Kriegeskorte et al., 2006). Using searchlights reduces the number of features, which is a way to avoid over-training, meaning a high performance in the training data but a poor performance in independent test data. Additionally, search lights provide information about the anatomical location of useful information within the brain. Conducting multiple, equivalent analyses, the search light is moved throughout the gray matter. We hypothesized that the highest accuracy will be obtained in regions of the value network (Ripke et al., 2012).

For our analysis, we used previously acquired data (Ripke et al., 2012). As a reference point, we initially evaluated the accuracy achievable using the offer details (amount and delay) and the behavioral models. We also evaluated the prediction performance of two SVMs (see Methods) using only these two offer features. Then, we evaluated the SVM classifier (see Methods for details) using fMRI data. Because the available data does not contain enough trials for classifier training at the indifference point only, we used all trials for this analysis. Thus, the obtained BOLD-based accuracies should reflect an upper limit for potential future studies with a focus on the presumably harder problem of predicting indifferent trials. To validate our methodology, we also conducted a performance analysis of the classifier for ‘left’ and ‘right’ motor responses and simulated to-be-expected accuracies assuming signals (features) that exhibit a realistic correlation with subjective value.

## 2 Methods

### 2.1 Data

For our analysis, we reused ITeCh data acquired within the project “The adolescent brain”, which has been funded by the German Federal Ministry of Education and Research (BMBF), and was previously published with a different emphasis using a conventional univariate general linear model (GLM) approach by Ripke et al. (2012). Within this projects, adolescents were studied at three different ages (14, 16, and 18). In each session, 90 IteCh trials were presented. Only the data of those subjects that participated in all three sessions was under consideration for this analysis. Of these 151 subjects, seven subjects were excluded because a mental disorder was diagnosed, 17 because they missed more than 10 trials, one because of implausible questionnaire responses and suspected lack of motivation, and five due to corrupted data. The remaining 121 subjects were randomly divided into a development set of 95 subjects and a validation set of 26 subjects.

The development set was used to fine tune our classification approach, while the validation set was used only a single time to validate our findings on independent data. Table 1 contains detailed information about the groups.

**Table 1:** Gender and mean age with standard deviation for the development and evaluation set of subjects.

| Set | N | Gender |  | Age Ses 1 |  | Age Ses 2 |  | Age Ses 3 |  |
| --- | --- | --- | --- | --- | --- | --- | --- | --- | --- |
|  |  | Male | Female | Mean | Std | Mean | Std | Mean | Std |
| Development | (95) | 51 | 44 | 14.53 | 0.32 | 16.46 | 0.33 | 18.51 | 0.38 |
| Validation | (26) | 12 | 14 | 14.48 | 0.38 | 16.43 | 0.42 | 18.53 | 0.45 |

Since the differences in the subjective value-related fMRI regression coefficient within the same subject at different ages is about as big as the differences between subjects (Fröhner et al., 2019), we decided to treat each session as independent.

All analyses presented were done within one session, i.e. every classifier was trained with a part of the data of one session, and tested with the remaining data of that session (see below).

### 2.2 Task

The task and the associated methodology has been described previously in (Ripke et al., 2012). Before the first fMRI session, a prior for the k and *β* parameters (see Sec. 2.2.3) was obtained for each subject to adjust the offers in the subsequent fMRI session such that the proportion of immediate and delayed choices would be approximately equal. For the same individual, the immediate amount was constant and know to the subject from the start of the experiment.

Ninety offers were presented in one fMRI session. At the beginning of each trial (see Fig. 1), subjects saw an amount of money and the associated delay for 2s. No immediate offer was displayed since it did not change within one individual. Then a fixation cross was shown for 6s, followed by 2s for the subject response and choice feedback. Subjects had to indicate their preference by pressing a button either with their left or right index finger. The motor response was cued and randomized for the responding hand with an exclamation mark that would indicate on which side the button had to be pressed for the later reward. Directly after their response, subjects received feedback in accordance with their choice (either the immediate or later reward). This was followed by a jittered inter-trial-interval of in average 7s resulting in 25 min. for one task run.

**Figure 1:**
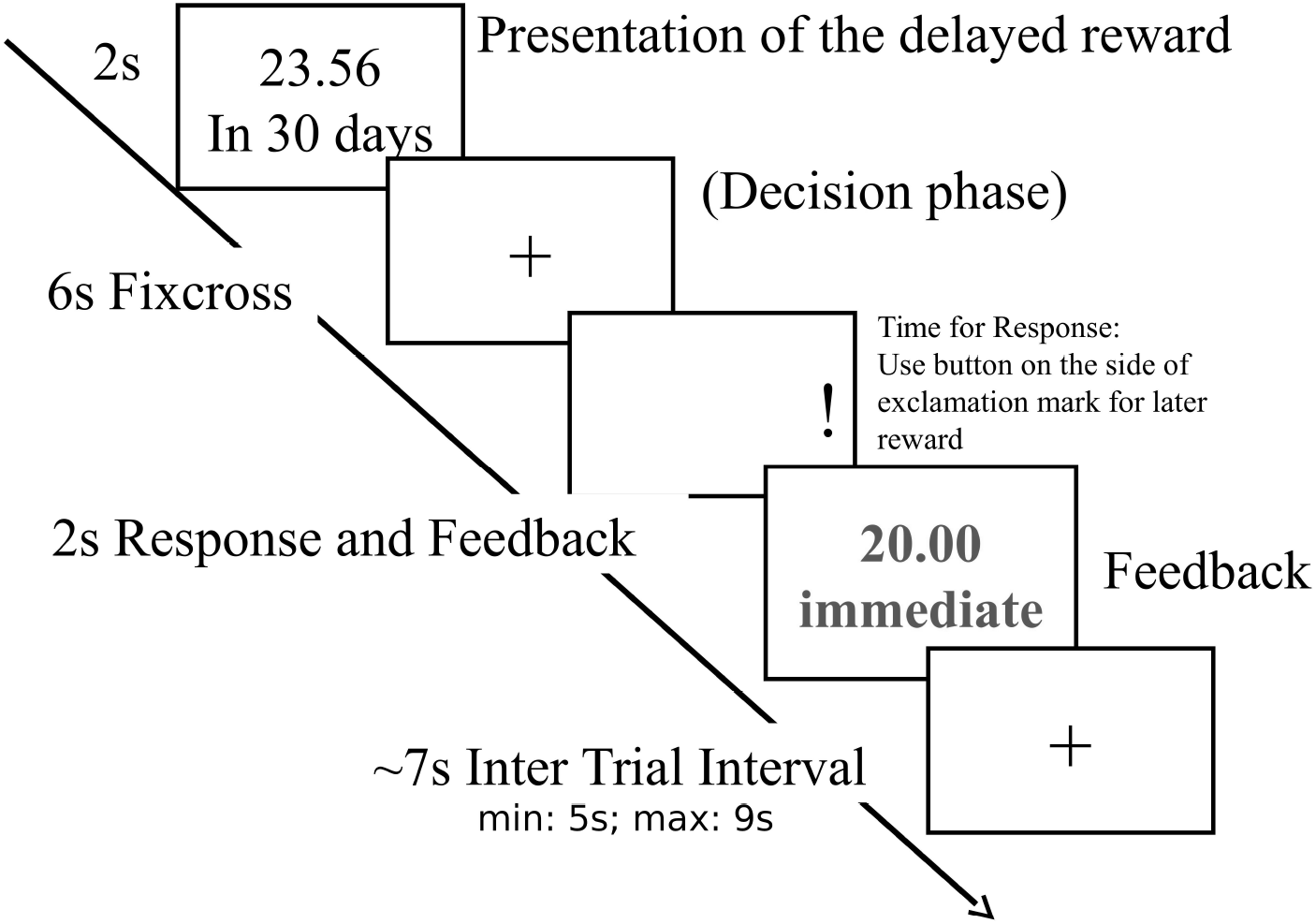
Illustration of the ITeCh task that was employed.

Using the amount and delay information in each trial and refitted *k* and *β* values based on subject choices, we computed the choice probability for each offer. We found that in 53% of the trials the delayed offer should be preferred and in 47% the immediate amount, respectively. 9% of all trials had a choice probability between 40% and 60% and were considered “hard” trials.

#### 2.2.1 MRI Acquisition Parameters and Preprocessing

Scanning was performed with a 3 T whole-body MR tomograph (Magnetom TRIO Tim, Siemens, Erlangen, Germany) equipped with a standard 12-channel head coil. For functional imaging, an echo planar imaging (EPI) sequence was used [repetition time (TR): 2410 ms; echo time (TE): 25 ms; flip angle: 80° bandwidth: 2112 Hz/pix]. Forty-two transverse slices were acquired, rotated 30° towards coronal from to the anterior commissure–posterior commissure line, with a thickness of 2 mm (1 mm gap), a field of view (FOV) of 192 × 192 *mm*^2^ and a matrix size of 64 × 64 pixels, resulting in a voxel size of 3 × 3 × 3 *mm*^3^. A structural 3D T1-weighted magnetisation-prepared rapid gradient echo (MPRAGE) image was also acquired (TR: 1900 ms, TE: 2.26 ms, FOV: 256 × 256 *mm*^2^, 176 slices, 1 × 1 × 1 *mm*^3^ voxel size, flip angle: 9°). Images were presented via NNL goggles (Nordic Neurolab, Bergen, Norway). Task presentation and recording of the behavioral responses was performed using Presentations software (version 11.1, Neurobehavioral Systems, Inc., Albany, CA). (Ripke et al., 2012)

For preprocessing, we applied slice-time and motion correction and transformed the images to MNI-space using SPM12 (Wellcome Centre for Human Neuroimaging, London, UK). No spatial smoothing was employed to maintain within-subject, multivariate patterns.

#### 2.2.2 Choice Prediction and Performance Evaluation

Throughout this paper, we are using within-subject 5-fold cross validation. This means that we divided the valid trials of one subject into 5 sets, with an, as much as possible, equal label distribution. We then merged 4 parts into the training set, and used the last remaining part as the test set. This procedure was repeated until every part was used as test set once, and the 5 resulting performance measures were averaged.

The most commonly quoted and most intuitive performance measure is accuracy or the percentage of correct predictions. The accuracy, however, can be a misleading statistic. If a data set has an unbalanced label distribution, the chance-level may be considered random guessing with a choice proportion as given in the training data resulting, which results in an accuracy larger than 50%. Our data sets have a number of different label distributions. Therefore, we computed Cohen’s *κ*, which normalizes for the label-distribution:

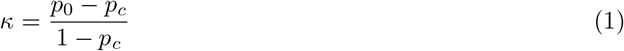

where *p*_0_ is the observed accuracy and *p*_*c*_ is the expected chance-level:

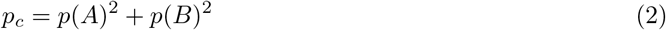

where *p*(*A*) is the choice probability of one class, and *p*(*B*) the choice probability of the other class based on the proportion of actually made choices.

Accuracies achieved by a classifier that only performs on the level of picking according to the label distribution will always produce a *κ* of zero. *κ* > 0 demonstrates performance above this chance level, and a *κ* = 1 a perfect prediction. *κ* < 0 means performance below chance level. While *κ* is technically a more appropriate parameter to quantify predictive performance, accuracy is easier to grasp and a more intuitive concept for readers that are unfamiliar with *κ*. Thus, we will provide a *κ*-equivalent accuracy Acc_50_ = 0.5 + 0.5 × *κ* assuming a 50:50 label distribution.

We had to exclude three runs from this analysis because they only had four labels of one class, and every fold must have at least one label for each class.

#### 2.2.3 The Behavioral Models

To put the fMRI-based prediction described below into perspective, we initially evaluated the performance of models that use only the offer details and subject behavior in the training set for prediction. We evaluated two models of this type (the Softmax and the Power model) and SVMs with a linear and Gaussian kernel using the delay and amount information as classification features.

The hyperbolic model computes a subjective value *V* given an amount *A*, a delay *D*, and the discounting parameter *k*. The formula is

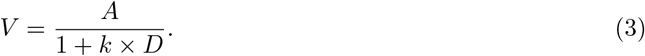

The parameter *k* is used to fit the model to subjects. Note that, independent of *k*, *V* = *A* when *D* = 0 (Fig. 2 a)).

We use the Bayesian framework by Pooseh et al. (2017) to fit the *k* and *β* parameters.

The difference between the two hyperbolic, behavioral models was the function that was used to calculate the probability for a particular decision given a particular subjective value for the delayed amount. The Softmax model uses the following function:

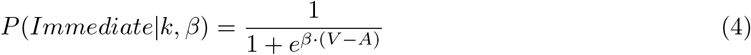

and

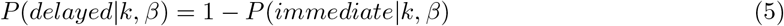

where *β* can be interpreted as a measure of consistency. For an intuitive explanation, see Fig. 2 b).

The Power-model uses:

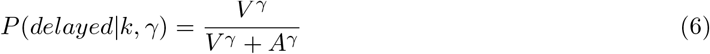

Here γ fills a similar role as *β* in Eq. 4.

**Figure 2.**
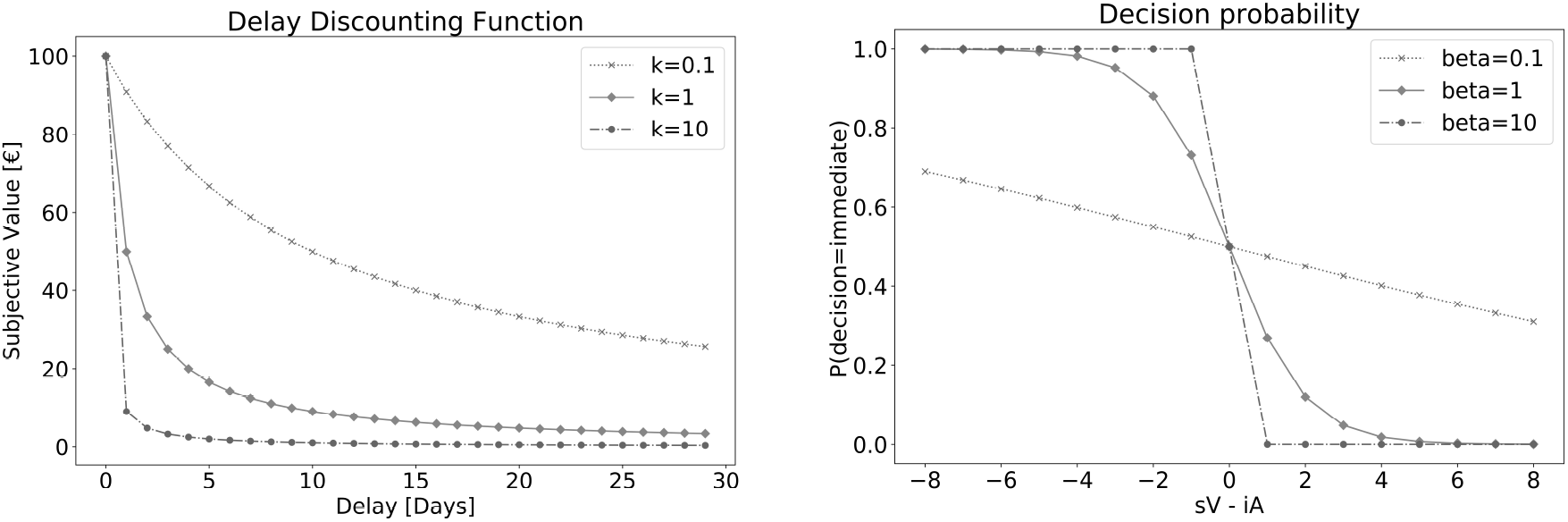
(a) Illustration of the hyperbolic delay discounting model. Subjective value of an amount is plotted over a delay of 30 days for 3 different values of *k*. As can be seen, larger *k*-values indicate stronger discounting. (b) The Softmax decision likelihood function (Eq. 4) for 3 different values of *β* as a function of (subjective value - immediate amount). For negative differences, it is more likely to choose the immediate offer, for positive ones, the delayed offer becomes more likely. It becomes more and more unlikely, that a subject will chose the immediate offer as difference between the subjective value and the immediate amount increases, i.e. the more desirable the delayed offer appears. The larger the *β*-value, the more this function approaches a step function and consistency increases.

Using these models, we built classifiers, which used the training data (4 of 5 folds) to fit a value for *k* and *β* or γ using a Bayesian fitting procedure (Pooseh et al., 2017) and then makes a prediction for previously unseen data, by computing the subjective value for the delayed offer using the fitted *k* value. If that subjective value is greater than the amount for the immediate offer, the classifier predicts ‘delayed’ otherwise it predicts ‘immediate’. Thus, the decision function only had an influence at the stage of fitting *k*.

As a more data-driven approach, we also used SVMs with a linear and a Gaussian RBF (radial basis function) kernel for prediction with a 2-dimensional feature-vector consisting of

1. offered amount for the delayed option
2. delay in days.

A C-value (a regulatisation parameter of the SVM) of one was used for both kernels.

### 2.3 BOLD fMRI-based Prediction

To generate the input features for the BOLD-based classification, we fitted a GLM with an event regressor at the onset of the offer presentation for every single trial during a run using SPM12. The canonical hemodynamic response function and a high-pass filter with a cut-off frequency of 1/(256s) were employed. The result was an individual coefficient map for every trial, which represented the BOLD signal approximately 6s post onset. We extracted feature sets for cubical search lights (SL) of 6×6×6 voxels centered on gray matter voxels with a grid spacing of 2 voxels in every spatial dimension. This led to an overlap of 67% between two neighboring SLs. For every searchlight and subject, a 5-fold cross validation with a linear SVM (regularization parameter C=1) was uses to compute *κ*-values.

We interpolated the resulting statistical maps by assigning all gray matter voxels the mean value of their direct neighbors ignoring none-gray matter voxels. On the group level, we then used a one-sided t-test for *κ* > 0 to compute p-values. We are aware that a permutation-based computation of the null hypothesis would be theoretically more appropriate than a t-test. However, this approach requires immense computational efforts to compute null-hypotheses for each search light and run. To investigate the impact of this alternative statistical test, we conducted permutation tests for a few search lights and found that the p-values were all smaller compared to the ones produced by the t-test. Thus, we decided in favor of the conventional t-test, which also has been employed frequently by other authors (Loose et al., 2017). Subsequently, we applied an FWE-correction to correct for the multiple searchlights. Region-labels were generated using the Neuromorphometrics atlas provided with SPM12.

### 2.4 Motor Response Classification

To validate our classification approach, we implemented the same approach described above to classify the left versus the right response hand using appropriate labels, which were randomized via the exclamation mark in the display (see sec. 2.2).

### 2.5 Simulation of Performance based on Correlated Variables

The second sanity check we conducted was a simulation to get an impression of realistic performances assuming BOLD signals that are correlated with subjective value. We know that the data shows a univariate correlation with subjective value, as was reported by Ripke et al. (2015). For this purpose, we generated 1, 2, 4, 8, 16, 32, 68, 128 and 216 features (voxels with BOLD signals for each trial) for each of the 185 IteCh runs (95 subject x 3 sessions) that had a realistic correlation of *r*=0.1 (Pearson) with the set of subjective values.

To find this ‘realistic’ correlation value, we correlated the timeseries of means of GLM regression coefficients within cubic ROIs in the Ventral Striatum and the Anterior Cingulate as reported by Ripke et al. (2012). We used the MNI coordinates (9, 9, −3), (−9, 6, −3), and (−3, 51, 0) as centers for 5 × 5 × 5 voxel cubes and average the regression coefficients within it. This led to 3 time series per subject which we correlated with the subjects subjective values. Averaged over all subjects in the developement set the correlations for the ROIs were 0.074, 0.085 and 0.072. Additionally, for the same ROIs, we computed r-values from Ripke et. al’s t-maps from the contrast that compared trials with choices for the delayed reward with trials for the immediate reward. These aerage r-values were 0.045, 0.048 and 0.038. Therefore we selected a correlation of 0.1 as an optimistic, for the purpose of this simulation conservative choice for our simmulation.

To compute a data series Y with a desired correlation with another data series *X*_1_, Eq. 7 was used.

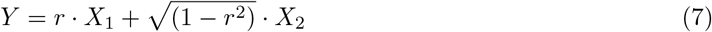

*X*_2_ is a set of numbers drawn from a gaussian distribution with a zero mean and a standard deviation of 1 with the same length as *X*_1_, and *r* is the desired Pearson correlation.

Our measures of interest were the resulting *κ*-values and statistical power as a function of the number of correlated BOLD signals. BOLD signals were not orthogonalized and would, therefore, show a marginal correlation with each other. Given that noise and signal in fMRI are spatially correlated, the simulated gain in accuracy by including additional voxels can be expected to be larger than for a real fMRI experiment as long as multi-variate, spatial correlations within the search light are not boosting classification performance. To compute power, we repeated the simulated experiment 300 times testing each time the average *κ* > 0 using a t-test. The multiple comparison issue was not addressed in this simulation.

## 3 Results

### 3.1 Behavioral Models

The behavioral models are summarized in Fig. 3. Both model-based classifiers and the linear SVM reached similar mean test accuracies around 90%. The Gaussian SVM performed worse, which could be a consequence of either the information being actually linear, so that there is no gain in the increased complexity of the Gaussian kernel, or the amount of training examples being too low for the increased complexity of the kernel. Still, all classifiers reached *κ*-values greater than 0.8. corresponding to accuracies Acc_50_ = 90%. A t-test for *kappa* > 0 resulted in *p* < 0.0001 for all approaches.

**Figure 3:**
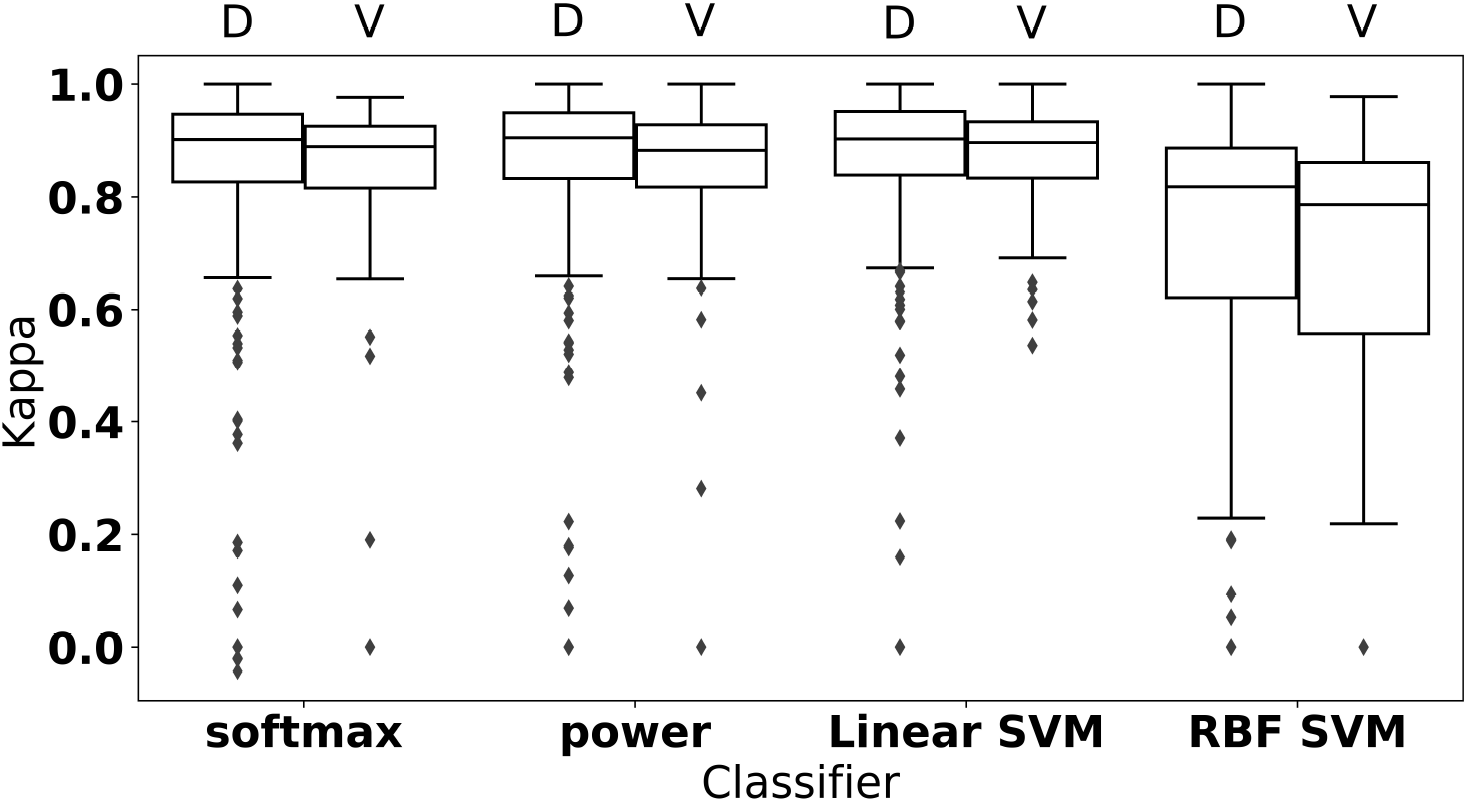
Box plots of mean test kappa values over sessions for the two behavior-based classifiers (Softmax and Power) and the two SVM approaches (linear and Gaussian (RBF) Kernal using the development set (D; N=285) and the validation set (V; N=78). A *κ*-value of 0.9 corresponds to Acc_50 = 95%.

### 3.2 fMRI Model

Fig. 4 shows the resulting t-map of the group level analysis of individual *κ* maps for the development set with 285 runs. Note that most of the gray matter has significant t-values, except for the motor area. The best performing searchlight is located at MNI=(−12,−64,56) near the left Precuneus with a performance of *κ* = 0.097 (*Acc*_50_ = 54.84%). The average over all significant voxels was 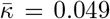 (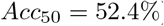)*STD*: 0.01

**Figure 4:**
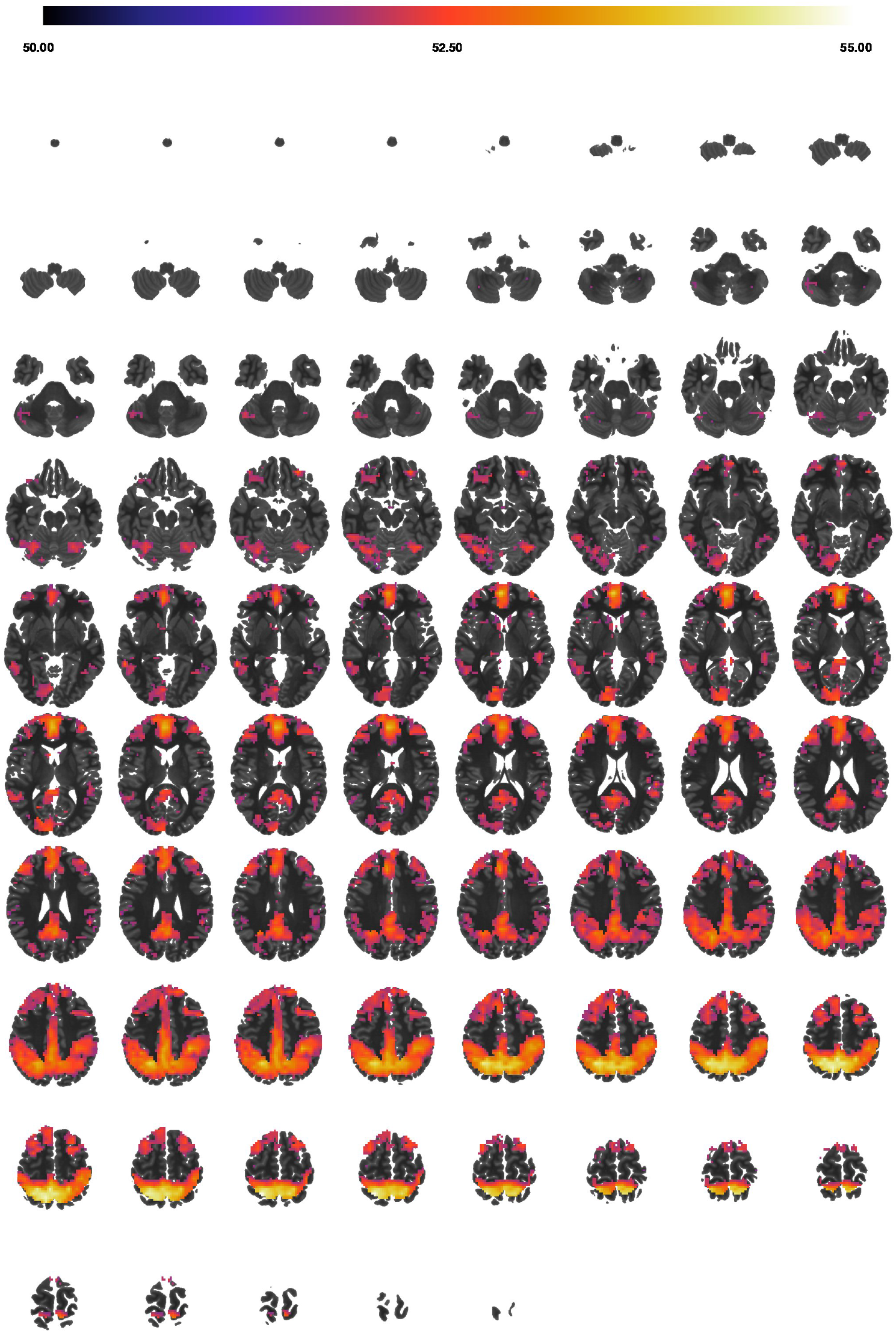
*Acc*_50_-map for the ITeCh choices in the development set thresholded at t = 4.77, which corresponds to an FWE-corrected p-value of *p* ≤ 0.05

Fig. 5 contains the resulting t-map from classifying the motor responses with summary statistics shown in Table 3. As to be expected, the primary sensorimotor areas show the most significant activation, with a peak *κ*-value of 0.12(*Acc*_50_ = 56.01%). The average over all significant voxels was *κ* = 0.062(*Acc*_50_ = 53.1%) STD: 0.019

**Figure 5:**
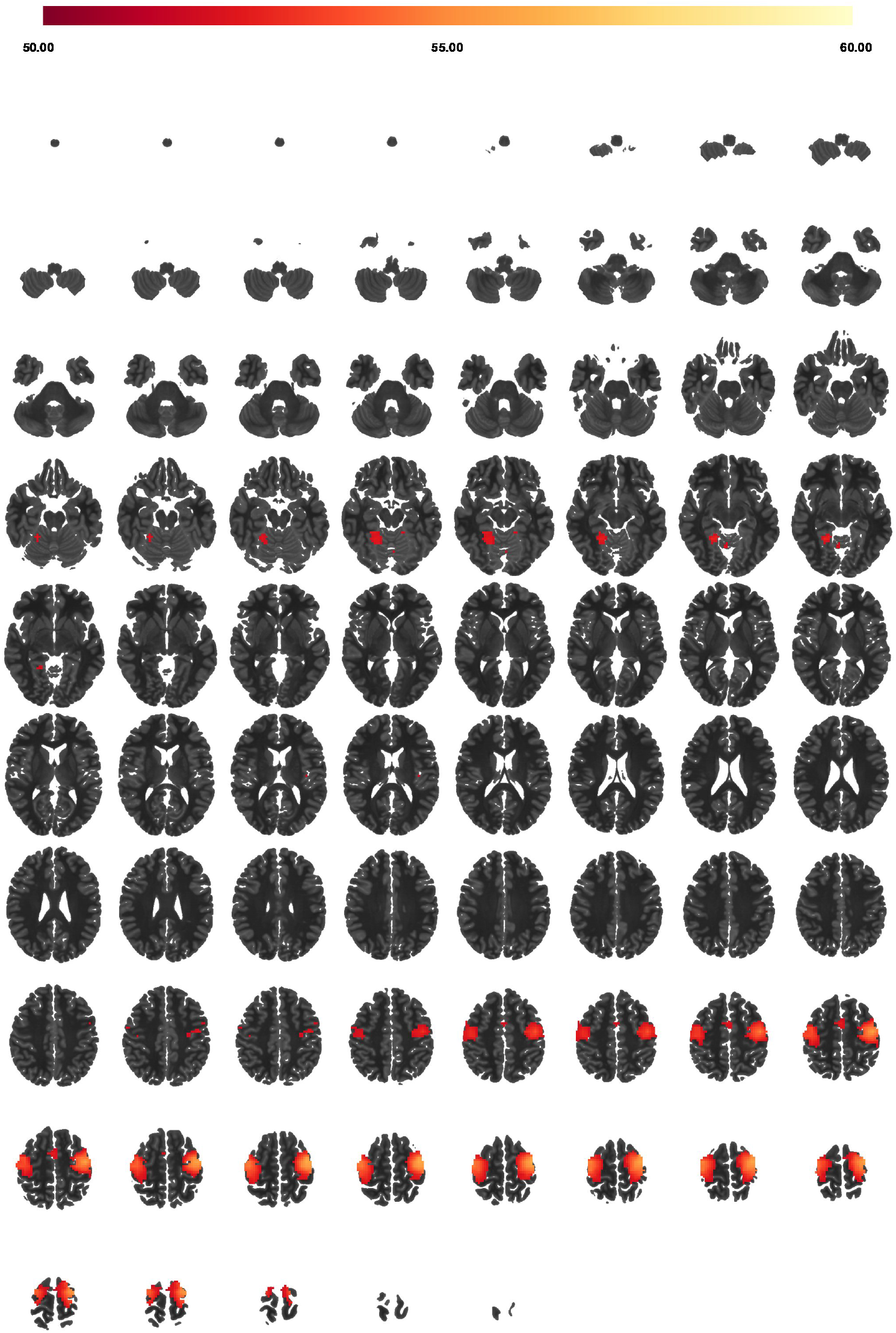
*Acc*_50_-map for the motor response classification in the development set thresholded at t = 4.77 as in Fig. 4

**Table 2:**
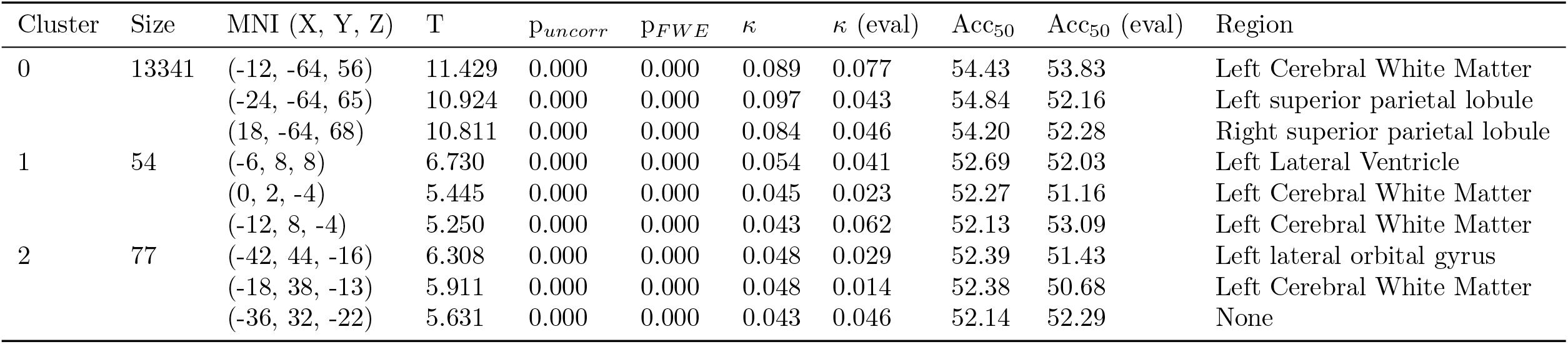
Summary statistics corresponding to Fig. 4 with 3 local maxima per cluster (with a minimum distance of 10 voxels) for all clusters with more than 50 voxels. The columns which contain the “(eval)” show the performance in the evaluation set at the same MNI coordinates.

| Cluster | Size | MNI (X, Y, Z) | T | $p_{uncorr}$ | $p_{FWE}$ | $\kappa$ | $\kappa$ (eval) | Acc<sub>50</sub> | Acc<sub>50</sub> (eval) | Region |
| --- | --- | --- | --- | --- | --- | --- | --- | --- | --- | --- |
| 0 | 13341 | (-12, -64, 56) | 11.429 | 0.000 | 0.000 | 0.089 | 0.077 | 54.43 | 53.83 | Left Cerebral White Matter |
|  |  | (-24, -64, 65) | 10.924 | 0.000 | 0.000 | 0.097 | 0.043 | 54.84 | 52.16 | Left superior parietal lobule |
|  |  | (18, -64, 68) | 10.811 | 0.000 | 0.000 | 0.084 | 0.046 | 54.20 | 52.28 | Right superior parietal lobule |
| 1 | 54 | (-6, 8, 8) | 6.730 | 0.000 | 0.000 | 0.054 | 0.041 | 52.69 | 52.03 | Left Lateral Ventricle |
|  |  | (0, 2, -4) | 5.445 | 0.000 | 0.000 | 0.045 | 0.023 | 52.27 | 51.16 | Left Cerebral White Matter |
|  |  | (-12, 8, -4) | 5.250 | 0.000 | 0.000 | 0.043 | 0.062 | 52.13 | 53.09 | Left Cerebral White Matter |
| 2 | 77 | (-42, 44, -16) | 6.308 | 0.000 | 0.000 | 0.048 | 0.029 | 52.39 | 51.43 | Left lateral orbital gyrus |
|  |  | (-18, 38, -13) | 5.911 | 0.000 | 0.000 | 0.048 | 0.014 | 52.38 | 50.68 | Left Cerebral White Matter |
|  |  | (-36, 32, -22) | 5.631 | 0.000 | 0.000 | 0.043 | 0.046 | 52.14 | 52.29 | None |

**Table 3:** Summary statistics when classifying the motor responses (’left’ versus ‘right’) with 3 local maxima per cluster (with a minimum distance of 10 voxels) for all clusters with more than 50 voxels. (PrCG: precentral gyrus, PoCG: postcentral gyrus, CWM: Cerebral White Matter). The columns which contain the “(eval)” show the performance in the evaluation set at the same MNI coordinate.

| Cluster | Size | MNI (X, Y, Z) | T | $p_{uncorr}$ | $p_{FWE}$ | $\kappa$ | $\kappa$ (eval) | Acc<sub>50</sub> | Acc<sub>50</sub> (eval) | Region |
| --- | --- | --- | --- | --- | --- | --- | --- | --- | --- | --- |
| 0 | 799 | (42, -16, 65) | 12.704 | 0.000 | 0.000 | 0.120 | 0.116 | 56.01 | 55.79 | Right precentral gyrus |
|  |  | (48, -16, 56) | 12.444 | 0.000 | 0.000 | 0.114 | 0.100 | 55.68 | 54.98 | Right postcentral gyrus |
|  |  | (33, -16, 71) | 11.816 | 0.000 | 0.000 | 0.108 | 0.108 | 55.42 | 55.40 | Right precentral gyrus |
| 1 | 587 | (-36, -16, 68) | 10.710 | 0.000 | 0.000 | 0.095 | 0.082 | 54.77 | 54.08 | Left precentral gyrus |
|  |  | (-42, -22, 62) | 10.477 | 0.000 | 0.000 | 0.092 | 0.070 | 54.62 | 53.49 | Left postcentral gyrus |
|  |  | (-36, -28, 68) | 10.027 | 0.000 | 0.000 | 0.090 | 0.093 | 54.50 | 54.66 | Left postcentral gyrus |
| 2 | 111 | (-24, -58, -10) | 6.814 | 0.000 | 0.000 | 0.056 | 0.013 | 52.78 | 50.63 | Left lingual gyrus |
|  |  | (-18, -64, -16) | 6.251 | 0.000 | 0.000 | 0.050 | 0.029 | 52.50 | 51.46 | Left Cerebellum Exterior |
|  |  | (-24, -52, -22) | 6.152 | 0.000 | 0.000 | 0.049 | 0.048 | 52.47 | 52.39 | Left Cerebellum Exterior |

### 3.3 Simulation Results

Fig. 6 summarizes the results of simulating a linear SVM classifier that used features (BOLD signals) correlated with subjective value. The power for our experiment to obtain a significant *κ*-value for our experiment is about 0.65 for a single feature and reaches 1 with 4 features. For 216 features, the average *κ* for all repetitions for all subjects reached 0.24. The peak *κ* of 0.097 from the fMRI-based IteCh choice prediction corresponds to approximately 30 nearly, independent features.

**Figure 6:**
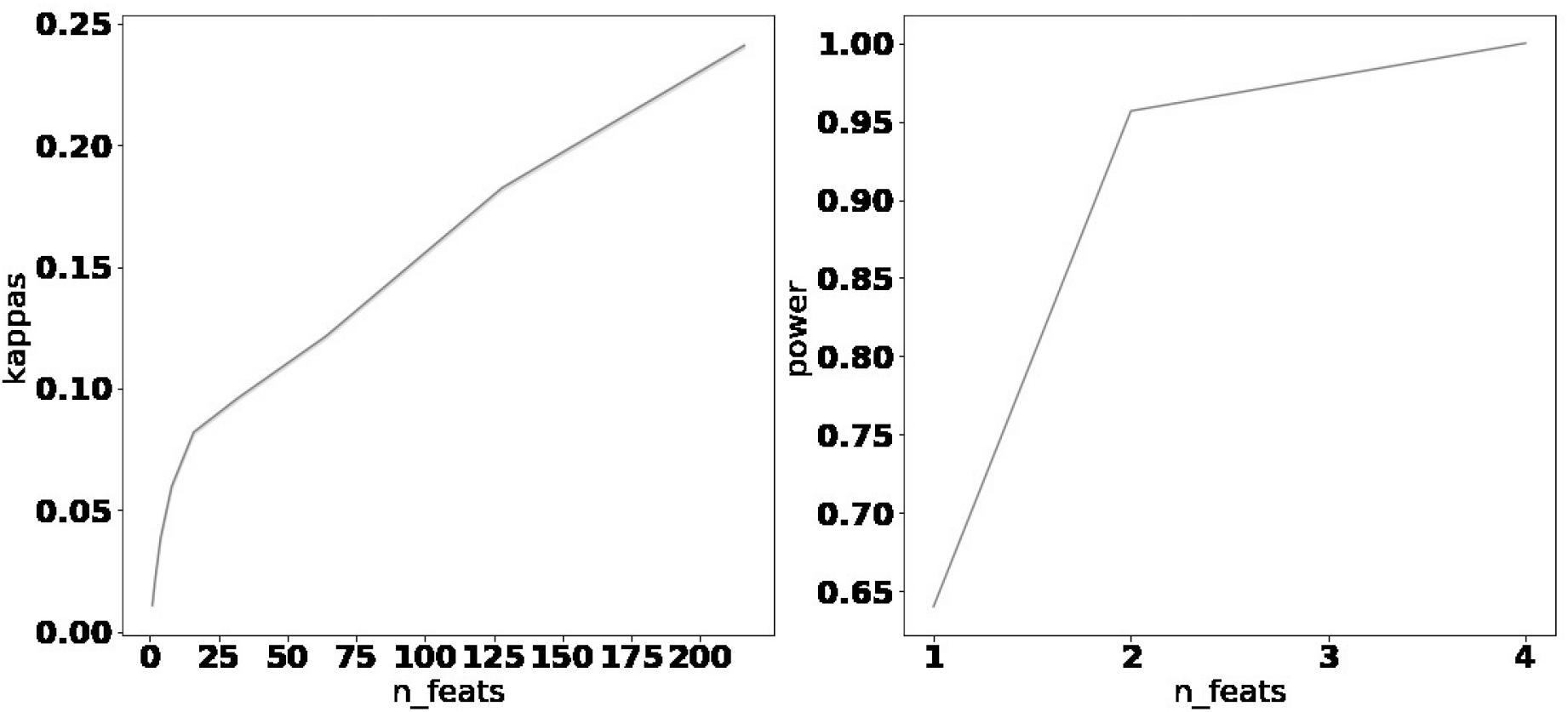
Results of simulating classifier performance based on a particular number of features (n feats) that are correlated with subjective value (*r* = 0.1) as computed for each trial in our experiment.

## 4 Discussion

This paper addresses two questions: 1. Does single trial BOLD activity allow to predict inter-temporal choices? And 2. Do single trial BOLD data provide additional insight into brain mechanisms and dynamic brain states that may be relevant to intertemporal choices? To this end, we compared prediction accuracies achievable by conventional behavior-based models with BOLD-based predictions using a search light approach. We analyzed previously acquired data with a proportion of hard trials, i.e. with choice probabilities between 40 and 60 %, of 9%. Our primary findings are that i. predictions based on offer details already reach very high accuracies of approximately 95% (*Acc*_50_) independent of the choice probability function (Softmax or Power) and also achievable by a conventional machine learning approach (linear SVM), and ii. BOLD-based classification accuracies using a search light approach only reach accuracies of Acc_50_ = 54.84%. This is clear evidence that fMRI data are far too noisy to capture the subjective value of ITeCh choices compared to behavior-based models at least for the mixture of hard and easy trials in our experiment and the classification approach employed by us. In fact, the fMRI features are also too noisy to gain any increase of prediction accuracy over the behavioral models through combining the offer amount and delay with the fMRI features. Our attempts (data not shown) resulted only in a reduction of the achievable accuracy, which is expected when adding features that are much noisier than existing features.

As stated in the introduction, it would be of particular interest to study trials that are hard, meaning by design not well predicted by the behavioral models. While we still believe that such an experiment would be of value, designing such an experiment will be challenging because it is reasonable to presume that the BOLD classification problem addressed in this paper with choices of mixed difficulty should be much easier than predicting hard choices only. But if choices with mixed difficulty are only predictable with 54% accuracy, it is likely that the achievable accuracy for hard choices will be even less. This leaves very little room to improve upon behavioral models. Exploring alternative classification methods that further optimize achievable accuracies, such as whole brain approaches, may improve upon this limitation. Secondly, we observed in pilot experiments that presenting only hard choices to subjects can lead to a change in subject behavior or strategy, e.g. they might decide to simply always take the immediate offer, without actually considering the delayed offer. This instability would be a confound for training a classifier.

We validated our methodology by classifying motor responses and showing that the highest accuracies for this task originate from sensorimotor regions as to be expected. It should also be noted that those same regions are not informative for classifying ITeCh choices, which is due to the randomization of motor responses. Somewhat surprising may be the low accuracy achieved for both the ITeCh choices and the motor response task of ‘only’ up to 56%. However, this is a consequence of low signal-to-noise ratio in fMRI experiments. We have to remember that we are analyzing single trials in this experiment, meaning a single button press. While motor activity is often used as a solid generator of BOLD signals, this is usually done in block designs with many movements over 10−30s.

Another way for us to put our findings into perspective are the simulations that we conducted assuming that a BOLD signal is present that tracks subjective value. The fMRI peak performance of *κ* = 0.097 is reached by the simulation with approximately 30 nearly independent, simulated features. It would be far too optimistic to assume the all 216 feature voxels can realistically be independently correlated with subjective value. In this light, the obtained *κ*-values appear reasonable to us. Potential spatial correlations exploitable by the classifier, however, are not considered here. The simulation also shows that obtaining significance even for low classification accuracies in a large sample like ours is easily possible even with only two features that correlate marginally (*r* = 0.1) with subjective value.

While the achieved BOLD-based accuracies are low, they are significant throughout most of the gray matter due to the large number of runs in the development set. The brain regions predictive for choice behavior demonstrate well-known networks that are involved in intertemporal decision-making (Weber and Huettel, 2008; Ripke et al., 2012; Peters et al., 2012; Kable and Glimcher, 2007) such as the valuation network (mPFC, PCC) and the cognitive control network that comprises mostly lateral regions of the frontal and parietal lobes (McClure et al., 2004; Peters and Büchel, 2011; Ripke et al., 2012).

In our results section, we only presented *κ*-values within a given search light averaged over all subjects. An obvious question is how well we could actually classify within a single subject if we were free to select the best possible search light. Unfortunately, this question cannot easily be answered because, if we simply choose the best search light (of 6000), we will find average accuracies of 84%, which are strongly inflated due to multiple testing issues (even stronger than to be expected by a binomial distribution). This accuracy collapses basically to chance level when evaluating it on previously unseen data, e.g. by using a nested cross validation, meaning that the selection process results in strong over-fitting. Preventing this problem is not trivial. An approach might be to use all search lights and make predictions by a majority vote. There are many approaches to address such a problem of feature selection and to optimize the achievable prediction accuracy further, for example through whole brain classifiers (Liu et al., 2015; Mohr et al., 2015). But this is beyond the scope of this paper.

Chen et al. (2018) published a similar study and reported much higher accuracies of up to 84% for individual decisions. We obtained only 55% accuracy. Even when averaging all trials per subject, we could only classify the decision-category (immediate or delayed) with an accuracy of up to 65% accuracy (average accuracy of the best searchlight). We can only speculate on the cause for this discrepancy based on the differences between the two studies. First, we used a searchlight, while Chen et al. used a whole-brain approach or particular regions of interest (ROI). We used within-session 5-fold cross validation in 285 sessions, while they used a between-subject approach (leave-one-subject-out and a 2-group approach). Also, the way they constructed the offers was different from ours, which might have led to easier decisions for the subjects and, in consequence, to higher accuracies in the classification process. Despite this discrepancy, we are confident that our accuracies represent realistic values for a typical IteCh task based on the motor response analysis and the conducted simulation.

## 5 Conclusion

We compared the performance of behavior-based and fMRI-based models for predicting ITeCh choices and showed that the behavioral models surpassed the fMRI model by far with accuracies of more than 90% in contrast to accuracies of 54% for the fMRI model. We also evaluated the classification performance of the fMRI model using a simulation, and arrived at the conclusion that the noise level in fMRI is strongly limiting the achievable prediction accuracy on single ITeCh trials. Tracking dynamic brain states of impulsivity will, therefore, be challenging. Nevertheless, informative brain regions can be identified with this approach that are consistent with the value and cognitive control networks. Further work to optimize classification accuracy particularly on trials hard to predict by behavioral models would be of interest.

